# Global immune fingerprinting in glioblastoma reveals immune-suppression signatures associated with prognosis

**DOI:** 10.1101/309807

**Authors:** Tyler J. Alban, Alvaro G. Alvarado, Mia D. Sorensen, Defne Bayik, Josephine Volovetz, Emily Serbinowski, Erin E. Mulkearns-Hubert, Maksim Sinyuk, James S. Hale, Giovana R. Onzi, Mary McGraw, Pengjing Huang, Matthew M. Grabowski, Connor A. Wathen, Tomas Radivoyevitch, Harley I. Kornblum, Bjarne W. Kristensen, Michael A. Vogelbaum, Justin D. Lathia

## Abstract

Glioblastoma (GBM) remains uniformly lethal, and, despite a large accumulation of immune cells in the microenvironment, there is limited anti-tumor immune response, even with newly developed immune checkpoint therapies. To overcome these challenges and enhance the efficacy of immunotherapies, a comprehensive understanding of the immune system in GBM and changes during disease progression is required. Here, we integrated multi-parameter flow cytometry and mass cytometry time of flight (CyTOF) analysis of patient blood to determine changes in the immune system among tumor types and over disease progression. Utilizing multi-parameter flow cytometry analysis in a cohort of over 250 patients with brain tumors ranging from benign to malignant primary and metastatic, we found that GBM patients had a significant elevation in myeloid-derived suppressor cells (MDSCs) in blood, but not immunosuppressive T regulatory cells. We validated these findings in GBM patient tissue and found that increased numbers of MDSCs in recurrent GBM portended poor prognosis. CyTOF analysis of peripheral blood from a cohort of newly diagnosed GBM patients revealed that reduction in MDSC frequency over time is accompanied by a concomitant increase in dendritic cells and natural killer cells. This reduced MDSC profile was present in GBM patients with extended survival and was similar to that of low-grade glioma (LGG) patients. Our findings provide a rationale for developing strategies to target MDSCs, which are elevated in GBM patients and predict poor prognosis, either by directly targeting or by shifting the immune profile to induce differentiation toward the immune profile of LGGs.

## Introduction

Glioblastoma (GBM) is the most prevalent primary malignant brain tumor and remains uniformly fatal despite aggressive therapies including surgery, radiation, and chemotherapy [1]. Currently, there is great interest in targeting the immune system to promote anti-tumor response as a new means of treating cancers, including GBM [2-4]. Despite the presence of potential anti-tumor effector cells within the microenvironment, GBM growth persists [5-14]. One possible explanation for the lack of effective anti-tumor immune response is the presence of a large number of immunosuppressive cells [7, 12-14]. This is likely due to a number of factors, including immune checkpoint signaling, T cell exhaustion, glucose depletion, hypoxia, and the presence of immunosuppressive cells, such as T regulatory cells, tolerogenic dendritic cells (DCs), and myeloid-derived suppressor cells (MDSCs) [12, 14, 15]. Another factor that contributes to the limited immune response could be that GBM has a low mutational load, which does not allow for the recognition and removal of cancer cells by the immune system [16]. All of these factors combined have led to testing of checkpoint inhibitors in clinical trials, which demonstrated that the antigen-specific T cell responses do not always correlate with tumor regression, suggesting that the immunosuppressive microenvironment limits the potential of T cell activation [17, 18]. While this amount of immunosuppression in GBM appears drastic, it is consistent with the immune-privileged nature of the brain and may thus be more difficult to reverse than with tumors in other locations [4, 19, 20].

Given these barriers to the use of immunotherapy approaches, identifying mechanisms of immunosuppression in GBM is an immediate priority. MDSCs are of particular interest given their previously identified role in GBM immunosuppression, immunotherapy response, and cancer progression [12, 13, 15, 21, 22]. A number of contact-dependent and contact-independent pathways have been described for MDSCs, which broadly inhibit T cell proliferation and activation [23]. MDSC production is induced following an inflammatory response to restore homeostasis [24]. However, it has been demonstrated that MDSCs are also increased in most, if not all, cancers in which they have been examined [25]. In addition, it is not surprising that MDSCs are also increased in GBM considering the dire consequences that large-scale inflammation in the brain could cause [26]. This opens up the possibility that while MDSCs could be induced by cancer cells to help evade immune recognition, they may also be recruited by healthy brain cells such as microglia and astrocytes to protect the brain from excessive inflammation [27]. We recently identified an interaction between MDSCs and GBM cancer stem cells via macrophage migration inhibitory factor (MIF) that leads to enhanced MDSC function, increased cytotoxic T cell infiltration, and could be targeted to reduce GBM growth [21]. However, due to the intricate and interconnected nature of the immune system, it is not clear how targeting a single immunosuppressive cell pathway would impact the function of the anti-tumoral immune system. Taken together this suggests that there is a need to delineate the complex nature of the GBM immune response.

The immune alterations in GBM have primarily been examined with targeted approaches such as immunohistochemical staining and flow cytometry, while RNA sequencing analysis (RNA-seq) remains the only unbiased approach [16, 28, 29]. GBM immunohistochemical analysis has been useful in identifying infiltrating macrophages/monocytes, T regulatory cells, and T cell dysfunction [6, 14, 30]. While immunohistochemical and immunofluorescent staining techniques are becoming more advanced, they fail to provide a general picture of the immune system within GBM [31]. Flow cytometry has similar pitfalls; although flow cytometry has been used successfully to identify immune cell infiltration and dysfunction in GBM, it is limited by the number of fluorescent markers that can be used due to compensation issues and overlap in fluorophore signals [32]. Finally, RNA-seq studies have been able to profile the immune response in GBM compared to other cancers, but this is a recent development in which the relative abundance of immune cell populations is determined using TCGA pan-cancer data [16]. These RNA-seq studies determined that intra-tumoral MDSCs, T regulatory cells, and effector memory CD4^+^ T cells are the most prevalent immune cell populations in GBM, with MDSCs enriched in more than 70% of patients with a low mutational burden [16]. While RNA-seq provides new insights into the immune landscape of GBM, these analyses have been performed on bulk tumors and are thus not an ideal way to examine multiple immune cell lineages to determine the composition of the immune microenvironment. An emerging technology with the potential to shed light on the immune microenvironment of many cancers is mass cytometry time of flight (CyTOF) [33-35]. CyTOF can identify immune cell response and differentiation, which can not be performed by traditional techniques [29]. This approach is currently being used in multiple cancers to examine the immune landscape of tumors in order to identify how to best enhance the anti-tumor immune response [29, 33, 34]. Here, we use a combination of approaches including flow cytometry, immunofluorescence, and CyTOF to identify an immunosuppressive phenotype with increased MDSCs and reduced anti-tumoral response in GBM patients with a poor prognosis compared to low-grade glioma (LGG) patients and GBM patients with a good prognosis.

## Results

### Flow cytometry analysis identifies increased MDSCs in GBM patients

To quantify immunosuppressive MDSCs in brain tumor patients, peripheral blood mononuclear cells (PBMCs) were isolated from patients undergoing surgical resection (**Figure 1A**). Samples were analyzed via flow cytometry using an MDSC-focused panel of antibodies against IBA1, HLA-DR, CD14, CD15, and CD33 (**Supplemental Figure 1A, C**). For comparison, a separate T cell-focused panel containing antibodies against CD3, CD4, CD25, CD8, CD107a, and CD127 was used to quantify cytotoxic T cells and T regulatory cells (Tregs) (**Supplemental Figure 1B, D, E**). The patient cohort for these studies utilized a total of 259 patients, who were subdivided into the categories benign, non-glial malignancy, and glial malignancy (grade I/II, grade III, grade IV (GBM)) (**Figure 1B**, with further final diagnoses for each group elaborated on in **Supplemental Figure 2**). When MDSC levels, as determined by the percent of HLA-DR^-/low^/CD33^+^/IBA1^+^ cells of the total live cells, were compared across groups, we observed that benign samples had a lower percentage of MDSCs compared to non-glial malignancies and grade IV glioma samples but not grade I/II or III glioma samples (**Figure 1C**). Additionally, non-glial malignancies had increased MDSCs compared to grade I/II tumors but not grade III or IV glioma, suggesting that MDSCs may be a possible marker of malignancy in brain tumor patients (**Figure 1C**). A direct comparison among glial malignancies within the categories of grade I/II, III and IV revealed that grade I/II tumors had significantly reduced MDSCs compared to grade IV samples, confirming results found by others (**Figure 1 C**) [15]. To determine whether other immunosuppressive cell types in circulation are also increased with malignancy, we assessed Tregs as the percentage of CD3^+^/CD4^+^/CD8^-^/CD127^-^/CD25^+^ cells (**Figure 1D, Supplemental Figure 1E**). No statistical difference was identified for Tregs among the categories of benign tumors, non-glial malignancies, glial malignancies, or other category. Returning to MDSCs, univariate analysis of dependence of MDSC level on age, sex, grade, isocitrate dehydrogenase 1 (IDH1) mutation status, O^6^-methylguanine–DNA methyltransferase (MGMT) status, and chronic steroid use prior to surgery (**Figure 1E**) yielded WHO grade as the most significant predictor of MDSC level (p=0.016). This remained marginally significant in bivariate models that controlled for the potentially confounding clinical variables age (0.016 to 0.076) and chronic steroid use (0.016 to 0.053), i.e. other variables with marginally significant univariate effects. These results demonstrate a relationship between circulating MDSCs and tumor grade but not between T regs and tumor grade.

**Figure 1.**
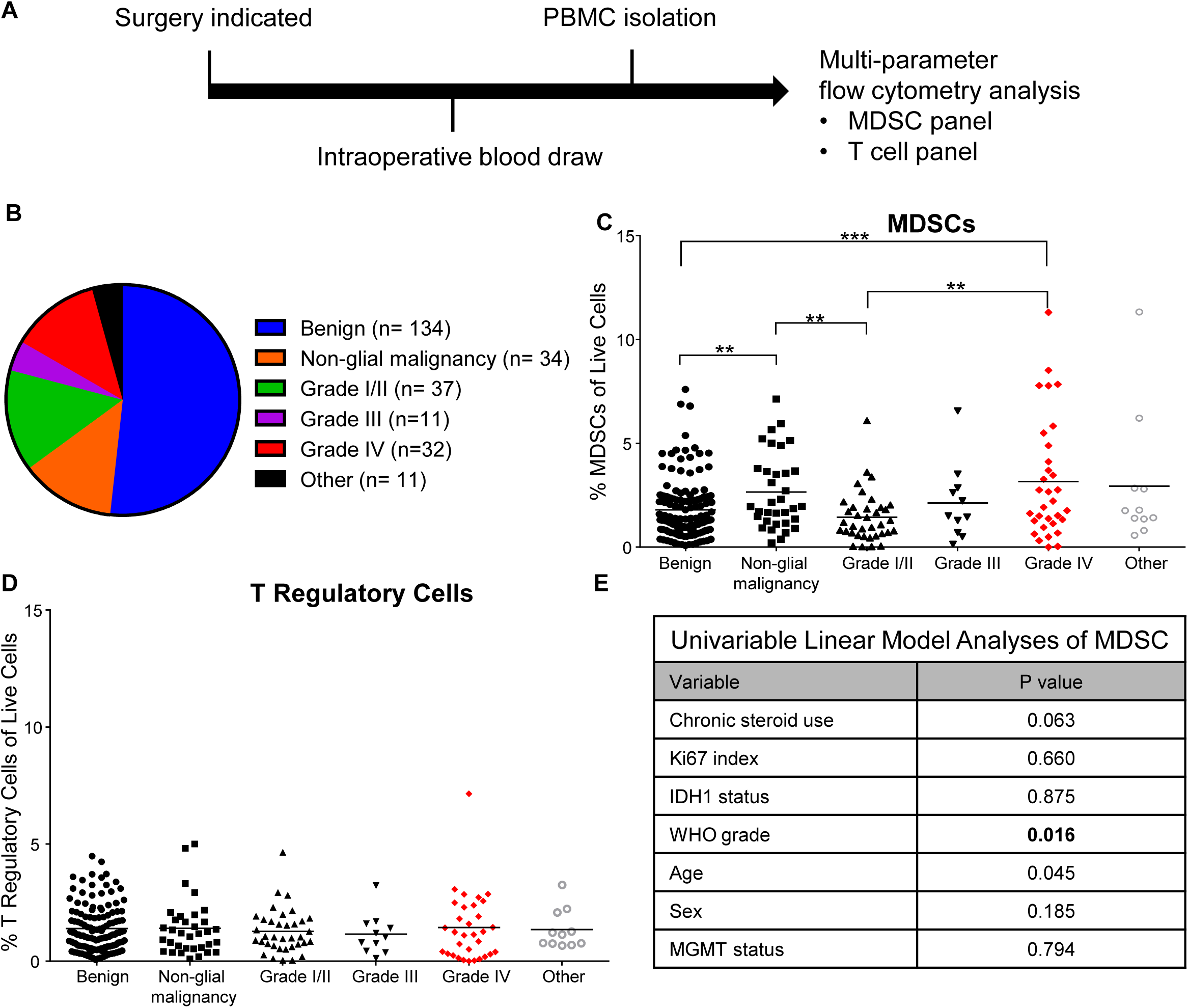
Multi-parameter flow cytometry analysis of blood samples from primary and secondary brain tumor patients reveals that GBM patients have increased immune-suppressive myeloid-derived suppressive cells. (**A**) Experimental design: patients entering the clinic for surgical resection were consented, and a blood sample was acquired intra-operatively. Subsequently, PBMCs were isolated via Ficoll-Paque™ gradient within 24 hours before being frozen in freezing media for future use. (**B**) Pie chart with the distribution of patient samples totaling 259 total patients analyzed. (**C**) Analysis of immune-suppressive M-MDSCs and T regulatory cells via multi-parameter flow cytometry analysis, where individual unpaired t tests were used to determine significant differences (* p≤0.05, **p<0.01, ***p<0.001). (**D**) Univariate linear model fits show that grade significantly associated with M-MDSC levels, while other clinical parameters were not significant (p≤0.05).

### Immunofluorescence staining of matched primary and recurrent GBM tumors identifies a correlation between M-MDSCs and survival

To validate our observation that circulating MDSCs were associated with increased malignancy, we utilized immunofluorescence analysis of MDSCs in paraffin-embedded matched primary and recurrent tumor samples from 22 GBM patients via antibody staining for CD33, IBA1, and HLA-DR (**Supplemental Figure 3, 4**). IBA1 was used in place of CD11b as CD11b has the ability to mark neutrophils and thus granulocytic MDSCs, while IBA1 should appear only on the M-MDSC compartment [36-38]. Within this cohort, patients were treated with a similar clinical paradigm (radiation and concomitant chemotherapy via the Stupp protocol [1]). Patients with high and low percentages of MDSCs were identified by the HLA-DR^low/negative^/IBA1^+^/CD33^+^ area relative to total tumor area using image analysis and grouped based on the MDSC signal in recurrent tumors. To determine whether an increase in MDSCs over the progression of tumors was associated with patient outcome, MDSC-high and MDSC-low groups were compared based on the median MDSC level, and we found that MDSC-high patients had a significantly reduced overall survival compared to MDSC-low patients (**Figure 2A**). This was not the case for overall myeloid cells as assessed by CD33 expression, as increased myeloid cell numbers were associated with increased survival (**Figure 2B**). Additional analysis of primary and recurrent resection samples for overall survival, time between first and second surgery, survival after the second surgery, and progression-free survival was performed for the correlation between each parameter (spearman r) and also via log-rank test (p-value) (**Figure 2C**). These analyses indicated that MDSC levels at primary resection were not predictive of prognosis but that MDSC levels during recurrence were informative for overall survival, time between first and second surgery, and survival after second surgery (**Figure 2C, Supplemental Figure 4**). These findings demonstrate that an increased infiltration of MDSCs portends poor prognosis, while infiltration of other subtypes of myeloid cells is beneficial.

**Figure 2.**
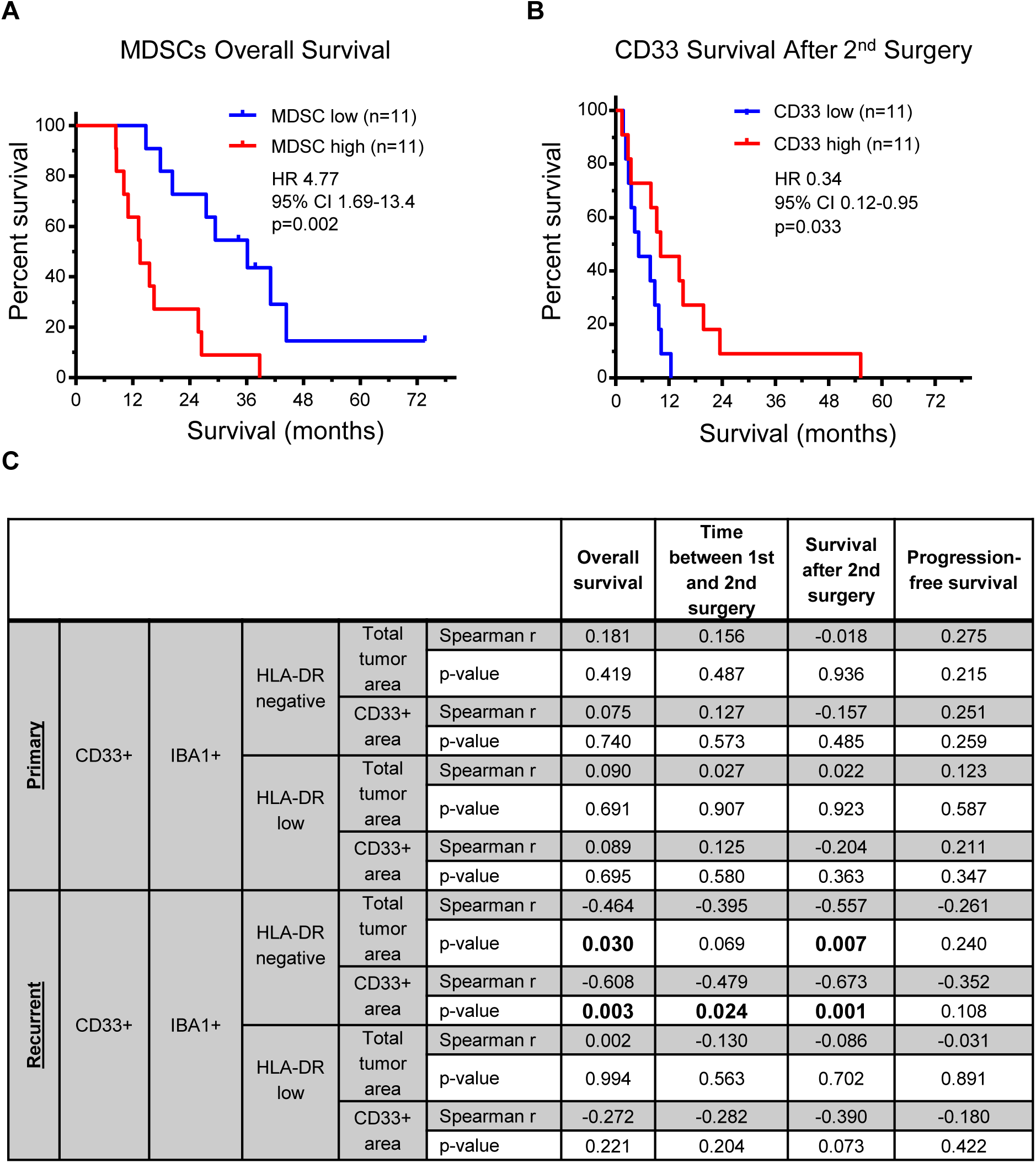
Immunofluorescence analysis of matched samples from primary and secondary resections from GBM patients identifies an associated between increased MDSCs and decreased survival. (**A**) Kaplan-Meier analysis of patients separated by median levels of MDSC signal in the CD33^+^ area demonstrates decreased survival. Statistical significance evaluated by log-rank analysis. (**B**) Kaplan-Meier analysis of patients divided by median CD33 levels identifies increased survival after 2^nd^ surgery using log-rank test (p=0.033). (**C**) Table of MDSCs separated by HLA-DR negative and low populations where correlation with survival, time between surgeries, survival after 2^nd^ surgery, and progression-free survival were analyzed (p<0.05 **Bolded**).

### Longitudinal study of GBM patients using an immune-fingerprinting approach via CyTOF reveals changes over disease progression

To determine how MDSCs and the overall immune system of GBM patients change during disease progression, samples from a cohort of 10 newly diagnosed GBM patients were analyzed via multi-parameter flow cytometry and CyTOF. Blood draws from these patients were obtained during surgery, two weeks post-surgery, and then every two months until the patient left the study or succumbed to disease (**Figure 3A**). Initially, all 10 patients were analyzed by flow cytometry using surface markers of MDSCs and T cells, described above. This analysis did not yield any significant trends in MDSCs or T cell populations (**Supplemental Figure 5, 6**). CyTOF was then used with a panel of 25 immune cell markers (**Supplemental Figure 7,8**) for a more in-depth analysis of how the immune system is altered during disease progression. For these analyses, three patients with a good prognosis (survival >600 days post-diagnosis, Patient 2= IDH mutant, Patient 3= IDH wild type, Patient 7= IDH wild type and three patients with a poor prognosis (survival <600 days post-diagnosis, Patient 4= IDH wild type, Patient 6= IDH wild type, Patient 9= IDH wild type) were selected. Samples were collected from each patient at three time points (baseline = intraoperative, 2 months post-surgery, final timepoint) and analyzed using multi-dimensional scaffolding analysis [39]. Unbiased clustering was performed to determine whether differences existed between the baseline values of patients and subsequent timepoints, and we observed that baseline samples grouped to one side of a multi-dimensional scaffolding (MDS) plot, indicating differences between the baseline and subsequent timepoints (**Figure 3B**). To identify cell type-specific clusters, a t-distributed stochastic neighbor embedding (tSNE) analysis was performed and, in an unbiased manner, identified 30 unique clusters of cells by taking 25 immune markers into account. The 30 clusters were then grouped into 12 immune cell types based on the histogram of marker expression within each cluster (**Figure 3C, Supplemental Figure 9**). Performing the same cluster analysis on each sample individually allowed visualization of how each immune cell cluster changed over time relative to the entire immune profile (**Figure 3D**). Integration of the CyTOF immune panel with multi-dimensional plotting identified immune cell shifts over time in a per-patient basis in an unbiased manner.

**Figure 3.**
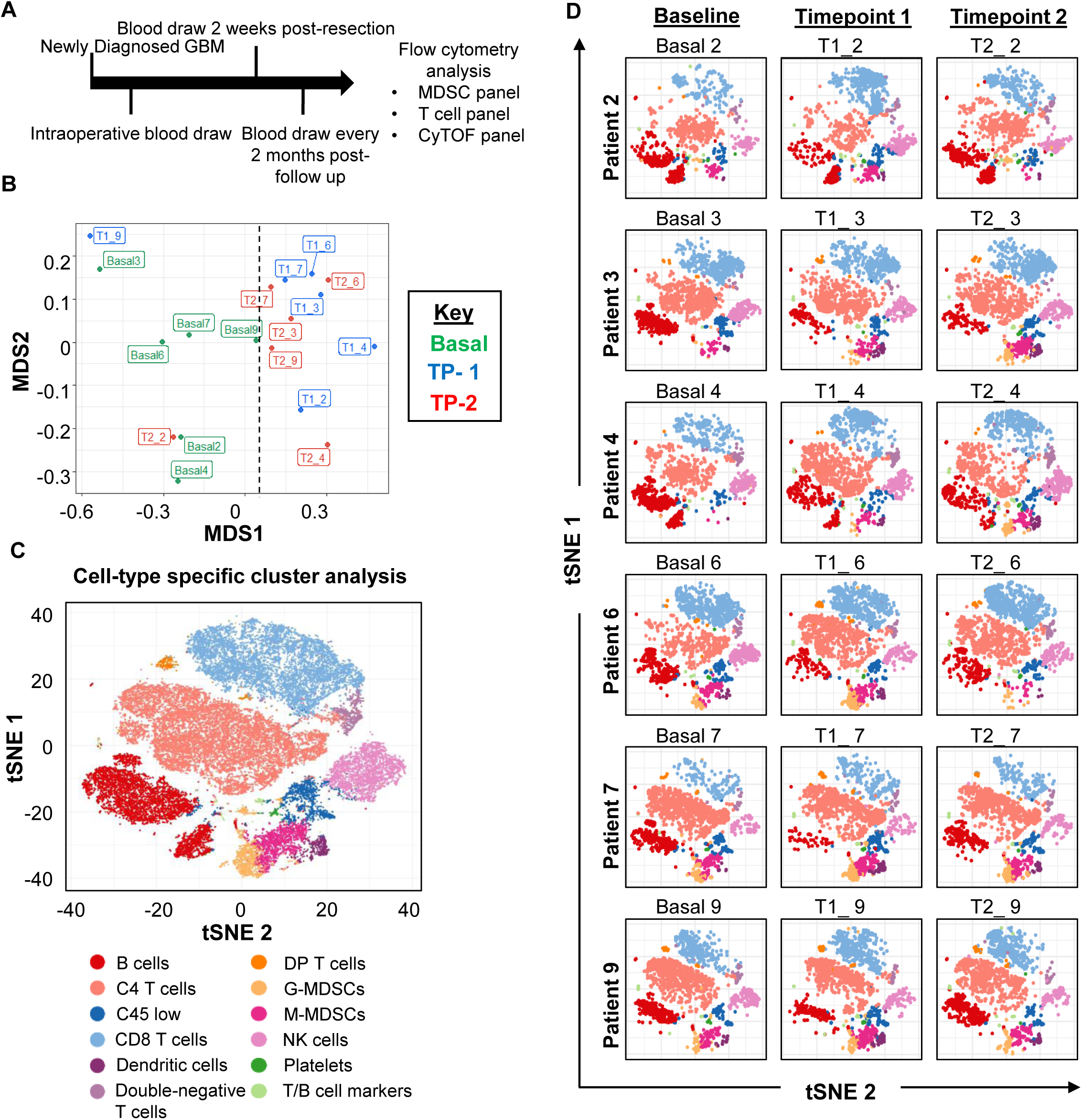
Mass cytometry analysis of GBM patients over time reveals immune shifts from baseline that are not common across all patients. (**A**) Schematic representation of the patient cohort consisting of 10 glioblastoma patients followed over time with blood collection and storage for analysis via multi-parameter flow cytometry and CyTOF. (**B**) Multi-dimensional scaffold plot representing 6 patients at three timepoints each (baseline, timepoint 1, and timepoint 2). The first number represents the timepoint, and the second represents the patient. Dotted line represents the division between baseline samples and later timepoint samples (**C**) t-distributed stochastic neighbor embedding (tSNE) plot identifies 30 unique populations that are color coded among the 6 patient samples across all timepoints, representing a total of 18 samples. (**D**) Individual tSNE plots of each sample demonstrate the quantity of each cell population by density of color-coded clusters over time.

### CyTOF analysis identifies changes in the immune system over time

To determine which immune cell populations changed over time, each population of immune cells was individually assessed. This analysis indicated that B cells, CD8^+^ T cells, DCs, and a mixed population of cells expressing a combination of B and T cell markers were significantly altered during disease progression (**Figure 4**). While B cells were significantly reduced compared to baseline, CD8^+^ T cells, DCs, and monocytic MDSCs (M-MDSCs) were significantly increased from baseline to 2-months post-surgery (**Figure 4**). While the increased CD8^+^ T cells and DCs are indicative of an anti-tumor immune response, there was also a reduction in B cells and an increase in immunosuppressive M-MDSCs and double-positive T cells, which are controversial and could be immunosuppressive or anti-tumor depending on the context [40]. Strong systemic immunosuppression was thus induced by the tumor. These data indicate that specific cell populations, including M-MDSCs, change during disease progression.

**Figure 4.**
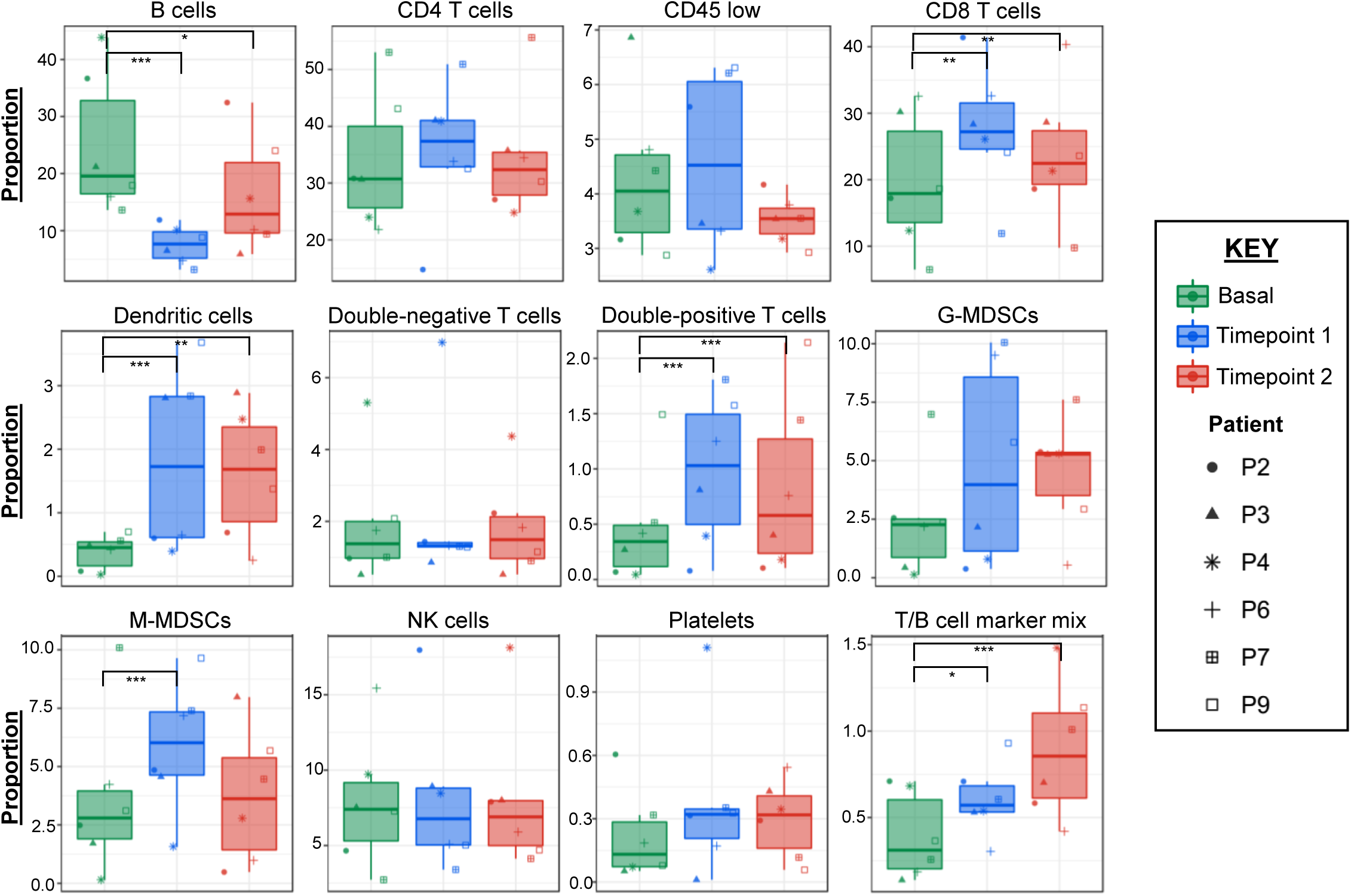
CyTOF identifies immune cell populations that are significantly altered during disease progression. Using 12 immune cell populations that were identified in an unbiased manner from baseline (green), timepoint 1 (blue), and timepoint 2 (red) samples for six newly diagnosed GBM patients were examined via student’s t test to compare baseline to timepoints 1 and 2. Each patient is indicated by the symbol identified in the KEY to the right. Statistics were determined by comparing baseline to each timepoint using Student’s t test *p<0.05, ** p<0.001, ***p<0.0001.

### CyTOF analysis between two patients with differing prognoses reveals changes in MDSC phenotype and overall immune cell differences

Based on our interest in MDSCs, their alterations during GBM progression, and their association with malignancy, we focused on MDSCs and the associated immune cell populations between patients with differing overall survival times. An in-depth CyTOF analysis of two patients from the longitudinal sample analysis was performed based on the correlation between their prognosis and changes in MDSC levels identified by flow cytometry (**Figure 5A, Supplemental Figure 7, 10**). Patient 2 had decreasing MDSCs over the course of the disease, had a favorable prognosis (survival >1,200 days), and was IDH1 mutant, while Patient 4 had increasing MDSCs over the course of the disease, a poor prognosis (survival of 583 days), and was IDH1 wild type. M-MDSC tSNE plots for Patients 2 and 4 show changes in the expression of markers in the CyTOF panel over time (**Figure 5B**). Corresponding heat maps of specific genes involved (**Figure 5C**) indicate that in both patients M-MDSCs increased Fc receptor/CD16 expression, and that Patient 4 had a 4-fold increase in CD61 that was not seen in Patient 2. CD61 signals by binding FGF1, which is known to be increased in GBM, and also enhances leukocyte rolling and adhesion, possibly indicating an increased ability to infiltrate the tumor and suppress the immune system within the tumor microenvironment [41, 42]. A FlowSOM analysis was performed on Patients 2 and 4 to determine the differences in immune response between the two patients, with an in-depth analysis of the markers expressed within each cluster and node, which were generated in an unbiased manner [43]. This analysis randomly clusters immune populations into 10 clusters containing unique nodes (indicated by spheres within the cluster), where the size of the node dictates the size of the population, and the internal pie charts indicate the expression of each marker within the node (**Figure 5D**). These results indicate that major differences exist within the anti-tumor immune response of patients with differing prognoses.

**Figure 5.**
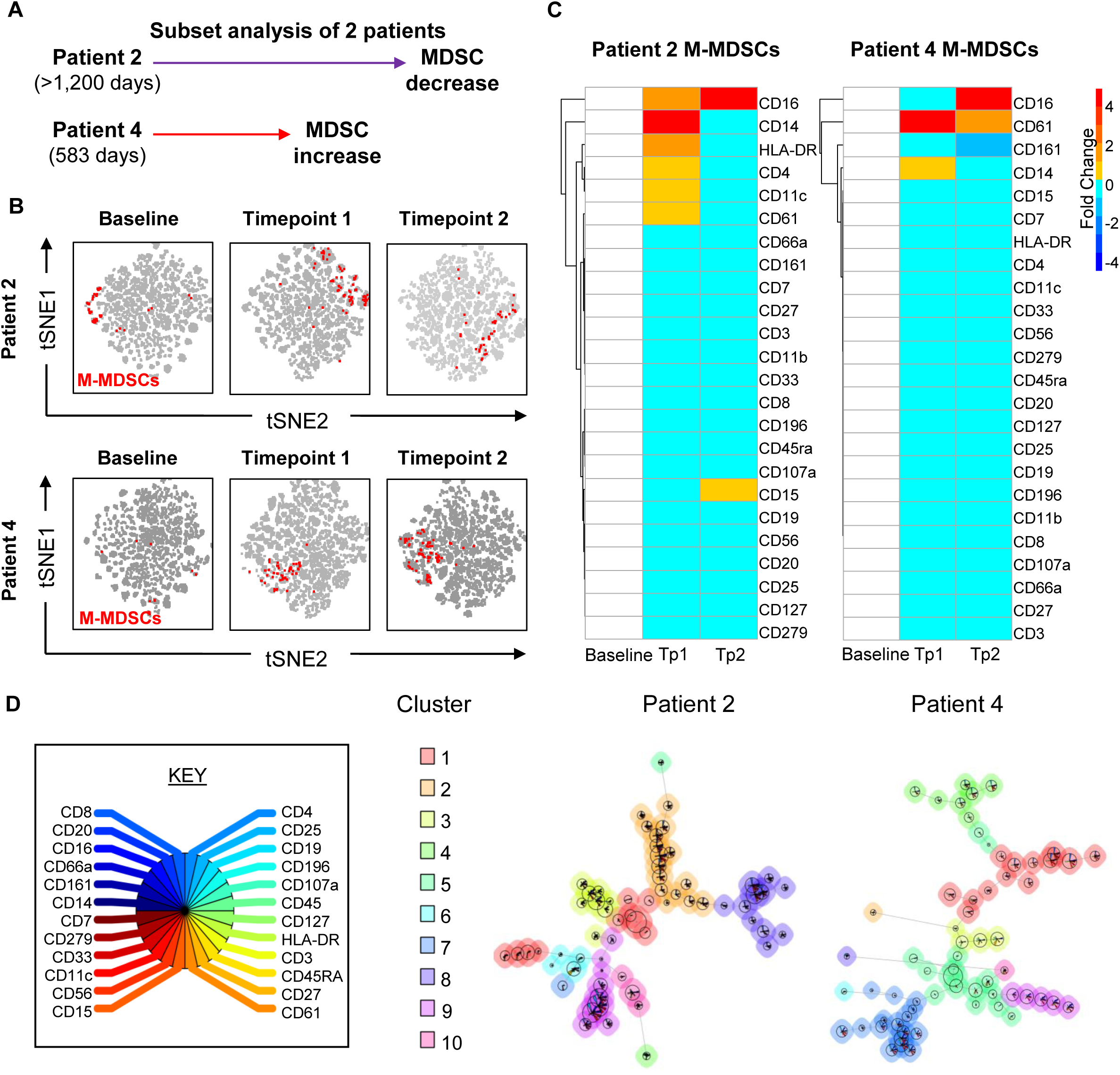
In-depth analysis of two patients with differing prognoses identifies shifts in MDSCs and other immune populations via FlowSOM. (**A**) Schematic representation of two patients used for in-depth manual gating analysis. Patients 2 and 4 were taken from the larger CyTOF study but were previously identified by multi-parameter flow cytometry as having decreasing and increasing MDSCs over time, respectively. Patient 2 had a survival greater than 1,000 days, while patients 4 had a survival of 583 days post-GBM diagnosis. (**B**) tSNE analysis of Patients 2 and 4 over time at baseline, timepoint 1 and timepoint 2, where manually gated MDSCs were overlaid and colored red. (**C**) MDSCs from Patients 2 and 4 were examined for fold change in markers from the CyTOF panel. (**D**) FlowSOM analysis of Patients 2 and 4 creates an unbiased clustering of 10 groups, with each node of the clusters identifying the size of the cell population and pie charts showing their expression of CyTOF markers.

### DCs and natural killer (NK) cells are increased in patients with a favorable prognosis

To gain a more in-depth appreciation of the changes in immune cell populations between Patients 2 and 4, 24 individual immune cell populations were manually gated (**Supplemental Figure 7**). We focused on MDSCs and antigen-presenting cells based on our earlier results (**Figure 3 and 4**) and NK cells based on their established role in the anti-tumoral immune response [12, 13, 21, 44, 45]. Compared to baseline, Patient 2 had a substantial increase in DCs at 2 months that was followed by an increase in anti-tumoral NK1 cells (CD45^+^, CD66a^-^, CD3^-^, CD19^-^, CD20^-^, CD14^-^, CD11c^-^, CD56^-^, CD16^+^) [46]. Additionally, M-MDSCs were reduced, as was previously identified by the multi-parameter flow cytometry staining, but an additional reduction in immunosuppressive NK2 cells (CD45^+^, CD66a^-^, CD3^-^, CD19^-^, CD20^-^, CD14^-^, CD11c^-^, CD56^+^, CD16^-^) was discovered using CyTOF (**Figure 6A**) [46]. Patient 4 did not show the same substantial increase in DCs or NK1 cells but instead had increased M-MDSCs and NK2 cells, which indicated an increase in immunouppression compared to Patient 2 (**Figure 6A**) [47, 48]. The reduction in MDSCs combined with increased DCs indicates the potential differentiation of MDSCs into DCs [49, 50]. To determine whether environmental conditions were favorable for MDSC differentiation, a 65-plex flow cytometry-based cytokine array was applied to serum samples obtained from Patients 2 and 4 at baseline, *t*_1_ (2 months) and *t*_2_ (last sampling time). This analysis revealed that FMS-related tyrosine kinase 3 ligand (FLT-3L) and granulocyte-macrophage colony-stimulating factor (GM-CSF), which are able to induce the differentiation of myeloid cells into DCs, were significantly increased in Patient 2 compared to Patient 4 (**Figure 6B**) [51]. Additional patients were examined via cytokine array, which revealed a clear signature of cytokine expression that differed between LGG and GBM patients (**Supplemental Figure 11**).

**Figure 6.**
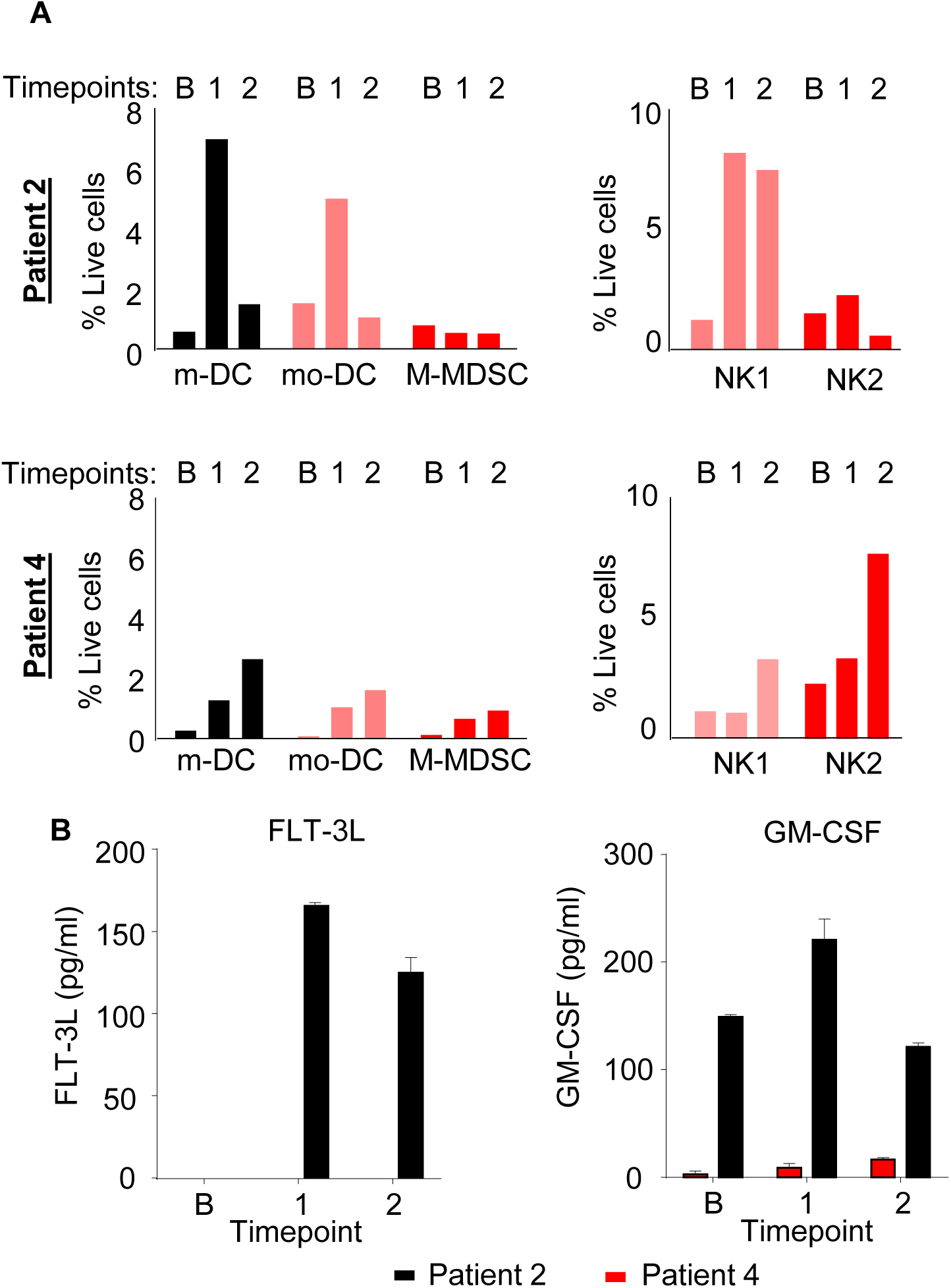
Dendritic cells and antigen-presenting cells are increased in a patient with a good prognosis. (**A**) Manual gating of dendritic cell populations, M-MDSCs, and NK cells from Patients 2 and 4 at baseline (B), timepoint 1 (1), and timepoint 2 (2), where B and 1 are at the same point in time post-diagnosis and 2 is the final time point collected. (**B**) Multi-parameter flow cytometry-based cytokine array where the serum levels (in pg/ml) of 65 cytokines were examined. FLT-3L and GM-CSF were increased in Patient 2 over time.

### CyTOF analysis of LGG patients reveals alterations in DCs and NK cells similar to those of a GBM patient with a favorable prognosis

Based on the distinct immune activation statuses found between two GBM patients (who each had a different survival status and IDH mutation profile), we compared six GBM patients to three LGG patients at diagnosis using samples on which we performed baseline CyTOF analyses (LGG1= IDH mutant, LGG2 = IDH wild type, LGG3 = IDH wild type). Although MDS revealed no clear difference between patients with GBM and LGG, tSNE analysis showed shifts in immune cell populations between GBM and LGG patients (**Supplemental Figure 12**). Further identification of the clusters within the tSNE and a quantification of immune cell changes revealed that the only significantly altered immune cell populations were DCs and NK cells, which were both higher in LGG patients than in GBM patients (**Figure 7A, B, Supplemental Figure 13, 14**). These results were consistent with our longitudinal study finding that a higher frequency of NK cells and DCs associates with a favorable prognosis and suggest that such GBM patients have an immune landscape similar to that of LGG patients.

**Figure 7.**
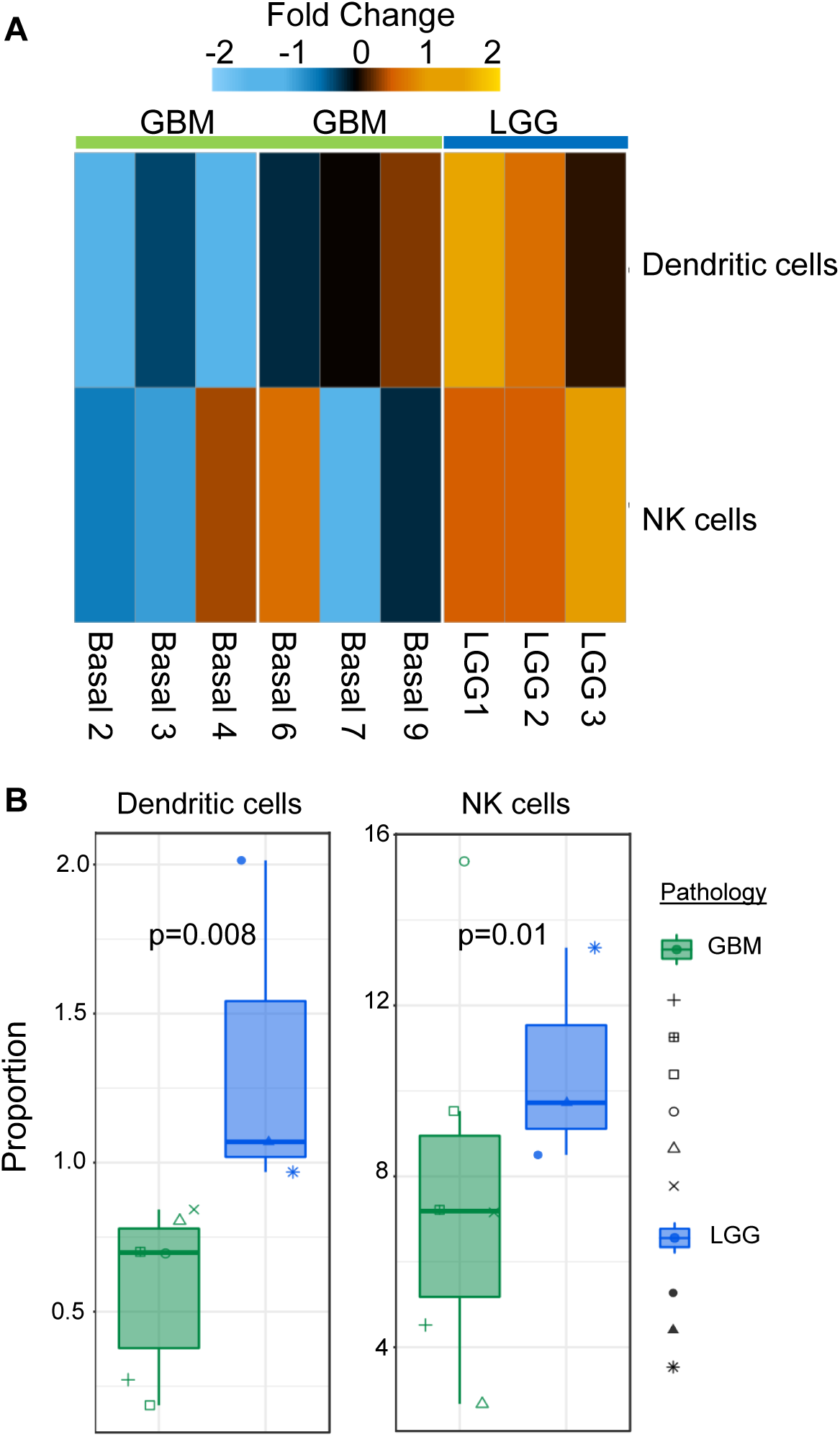
Compared to LGG patients GBM patients have reduced antigen-presenting cells and NK cells, which is indicative of a reduced anti-tumoral response. (**A**) Unbiased clustering of CyTOF data identifies NK cells and dendritic cells as different between patients with LGG and GBM at baseline as organized by hierarchical clustering. (**B**) Quantification of NK cells and dendritic cells in six GBM patients and three LGG patients at baseline using the t-test.

## Discussion

The correlations between peripheral anti-tumoral immune response and tumoral immune response have been of great interest; however, the identification of the peripheral immune status of GBM patients compared to that of patients with other types of brain tumors has not been comprehensively assessed. As the field of tumor immunotherapy progresses, it is vital to determine how the systemic immune response is altered under various tumor diagnoses, as past experiences have revealed that one drug does not work for all patients with the same disease, and it appears that immunotherapies are encountering a similar roadblock [52]. To identify new immunotherapeutic approaches or to enhance the efficacy of existing ones, we must first understand the immune landscape that is altered by the tumor and then ask how the drug of interest impacts that landscape.

Within GBM, patients have a skewed immune system with increased immunosuppression. However, studies typically focus on only one or two immune cell types of interest and do not examine the immune landscape as a whole or the immune response relative to other brain tumors. Here, we have developed a focused CyTOF panel to provide an understanding of the immune system as a whole and to predict how immune-modulating therapies may impact the anti-tumoral immune response of patients [53]. This understanding will aide in the investigation of future drugs in an unbiased manner by analyzing immune cell types implicated in immunosuppression and activation within GBM. We hypothesized that MDSCs are increased in GBM patients compared to patients with other types of brain tumors, based on their increased malignancy, and that the systemic immune response to GBM may differ over time among patients based on their prognosis and diagnosis. Our findings support this hypothesis and reveal that GBM patients with a more favorable prognosis exhibit decreased MDSCs and increased DCs, suggesting that MDSC differentiation is associated with an increase in immune activation and thus a decrease in GBM growth.

Through these studies, we found that immunosuppressive MDSCs are elevated in high-grade glial malignancies and in non-glial malignancies with brain metastases, while suppressive T cell populations were not altered as previously reported [28, 54, 55]. This is important as many therapeutic strategies currently under investigation for GBM aim to activate the immune system, as opposed to targeting the immunosuppressive cell types induced by the tumor [3]. While systemic immunosuppression was observed in these studies, we also observed immunosuppression intra-tumorally where MDSCs correlated with overall survival. This observation was made using matched primary and recurrent tumor-resection samples, where elevated levels of CD33^+^ myeloid cell infiltration correlated with a good prognosis, while infiltration of a specific subtype of myeloid cells, MDSCs, into the tumor microenvironment correlated with poor prognosis. These findings align with the genomic analysis of the immune landscape of IDH wild type and mutant gliomas previously identified [56]. Based on these findings, future studies could be performed to confirm the utility of MDSCs as a biomarker of disease malignancy and progression in brain tumor patients.

To gain an understanding of how patients’ immune systems change over time with disease progression, a CyTOF panel focused on 25 immune markers was developed that identified alterations in immune activation status and immunosuppression as a function of time. Specifically, while DCs and CD8^+^ T cells were increased over time, there was also a corresponding increase in immunosuppressive M-MDSCs and a decrease in B cells. The increase in DCs is of particular interest because it has been shown that DCs from the circulation are more effective at activating an anti-tumoral immune response than resident cells with MHC II such as microglia [57-59]. This phenomenon of immune recognition without an anti-tumoral immune response has also been observed in clinical trials of immunotherapies [59]. The associations identified between GBM patients and their status of immunosuppression with increasing MDSCs paves the way for future studies to combine anti-MDSC therapy with immune checkpoint therapies to enhance efficacy. Based on the differences noted between LGG patients and GBM patients by multi-parameter flow cytometry analysis, CyTOF was performed at baseline for LGG and GBM patients. DC and NK cell levels were higher in LGG patients compared to GBM patients, which could indicate that LGG patients are primed for an antigen response prior to surgery and are thus better able to mount an anti-tumor immune response to some degree. On the basis that MDSCs have the ability to differentiate into dendritic cells [23, 50], the data here suggest that low-grade tumors favor MDSCs maturing into DCs and that high-grade tumors favor MDSCs remaining as MDSCs. Future studies targeting differentiation pathways could enhance the anti-tumoral immune response and increase the efficacy of immune checkpoint therapies.

## Materials and Methods

### Study design

We sought to determine the relative frequency of MDSCs in GBM patients compared to patients with other primary and secondary malignant and benign brain tumors and, using CyTOF technology, to determine how the immune system of GBM patients is altered. Blood samples from a total of 260 patients were collected from brain tumor patients entering the Cleveland Clinic for treatment under Cleveland Clinic IRB 2559. Patients were grouped by their diagnoses into categories Benign, Non-glial malignancy, Grade I/II, Grade III, Grade IV, and Other as outlined in **Supplemental Figure 2**. Additionally, a cohort of 10 newly diagnosed GBM patients was enrolled in a blood collection study where blood samples were drawn every 2 months, with samples stored for general use by the Rose Ella Burkhardt Brain Tumor and Neuro-Oncology Center. Patient data was blinded from the researchers by the Rose Ella Burkhardt Brain Tumor and Neuro-Oncology Center through the generation of a de-identified numbering system. Multi-parameter flow cytometry and CyTOF panels were designed with MDSC and T cell populations in mind based on their relevance to GBM and previous identification within GBM (**Supplemental Figure 7**). Additionally, tumor tissue from 22 patients was retrospectively investigated from Odense University Hospital, Denmark. All patients were diagnosed with primary GBM between 2007 and 2015 and had not received any treatment prior to initial surgery. Following initial surgical resection, all patients received radiotherapy and chemotherapy. All patients experienced tumor recurrence within 31 months (mean progression-free survival: 13.3 months; range: 4.9 to 30.4 months), and the time period between initial and surgery resection was 15.2 months on average (range: 5.1-37.4 months **Supplemental Figure 4**). Four patients were diagnosed with recurrent GBM of the subtype gliosarcoma, the remaining 18 were diagnosed with recurrent GBM. All tissue samples were re-evaluated according to World Health Organization (WHO) guidelines 2016. Use of tissue was approved by the official Danish ethical review board (Named the Regional Scientific Ethical Committee of the Region of Southern Denmark), which approved the use of human glioma tissue (permission J. No. S-2011 0022). Use of the tissue was not prohibited by any of the patients according to the Danish Tissue Application Register.

### Flow cytometry

Peripheral blood samples were analyzed to determine MDSC and T cell populations in GBM patients over time as well as in low-grade glioma patients and was carried out in accordance with an approved Cleveland Clinic Foundation IRB protocol. Upon arrival, samples were processed through a Ficoll gradient in Ficoll-Paque PLUS and SepMate™ (Stem Cell Technologies) tubes before being suspended in freezing medium for storage. Samples were stained with live/dead UV stain (Invitrogen) and then blocked in FACS buffer (PBS, 2% BSA) containing FcR blocking reagent at 1:50 (Miltenyi) for 15 minutes. After live/dead staining and blocking, antibody cocktails (**Supplemental Figure 1**) were incubated with samples on ice for 25 minutes before being washed and suspended in FACS buffer. Cell populations were analyzed using an LSRFortessa (BD Biosciences), and populations were separated and quantified using FlowJo software (Tree Star Inc.). Gating methods for MDSCs were performed following standardized gating strategies previously described and outlined in **Supplemental Figure 1**, where MDSCs are marked by IBA1^+^, CD33^+^, HLA-DR-/low and can then be further subdivided into granulocytic MDSCs (CD15^+^) and monocytic MDSCs (CD14^+^) [60]. T regulatory cells were gated as CD3^+^, CD4^+^, CD25^+^, CD127^-^ as previously described [61]. CD8^+^ T cells were gated on CD3^+^, CD8^+^, CD4^-^, which were then determined to be activated by expression of the degranulation marker CD107a [62].

### Immunofluorescence

Fresh tissue biopsies were fixed in 4% neutral buffered formalin and embedded in paraffin. Sections (3 μm) were used for triple immunofluorescence staining, which was performed on a Dako Autostainer Universal Staining System (Dako, Glostrup, Denmark). Heat-induced epitope retrieval was performed in a buffer solution consisting of 10 mmol/L Tris base and 0.5 mmol/L EGTA, pH 9, followed by blocking of endogen peroxidases with hydrogen peroxide. Sections were then incubated for 60 min with a primary antibody against CD33 (NCL-L, Novocastra, Newcastle, UK, 1:200), and the antigen-antibody complex was detected using CSA II Biotin-free Tyramide Signal Amplification System kit (Dako) with fluorescein as the fluorochrome. After a second round of Heat-Induced Epitope Retrieval (HIER) followed by endogenous peroxidase quenching was performed, sections were incubated with an anti-HLA-DR antibody (CR3/43, Dako, 1:200) for 60 min, and Tyramide Amplification Signal Cyanine 5 (TSA-Cy5, Perkin Elmer, Waltham, Massachusetts, USA) was used as the detection system. Sections were then washed and incubated with an anti-IBA1 antibody (019-19741, 1:300, Wako Pure Chemical Industries, Osaka, Japan) for 60 min followed by detection with a goat anti-rabbit Alexa 350 secondary antibody (A-110461, 1:100, Invitrogen). Coverslips were mounted using VECTASHIELD Mounting Medium (VWR International, Radnor, Pennsylvania, USA). Omission of primary antibodies served as negative control. Fluorescent imaging and quantitation were carried out using the Visiopharm integrated microscope and software module (Visiopharm, Hoersholm, Denmark) consisting of a Leica DM6000B microscope connected to an Olympus DP72 1.4 Mega Pixel CCD camera (Olympus, Tokyo, Japan) using DAPI (Omega XF06, Omega Optical, Brattleboro, USA), FITC (Leica, Wetzlar, Germany) and cyanine-5 (Omega XF110-2) filters. Super images were acquired at 25x magnification using brightfield microscopy. Next, sampling regions were manually outlined. Sample images were collected using systematic uniform random (meander) sampling at 20x magnification with a minimum of five images per tumor. Images were reviewed to ensure that no artifacts or blurring were present. Images were then analyzed to quantify the amount of MDSCs in each tumor using a threshold-based algorithm developed in the Visiopharm software module. CD33 was used as an inclusion marker, and the algorithm was designed to identify the entire CD33^+^ area within the total tumor area. The CD33^+^ area was then subdivided into IBA1^+^ and IBA1^-^. The CD33^+^ IBA1^+^ area was then separated into three areas based on the intensity of HLA-DR staining: 1) CD33^+^ IBA1^+^ HLA-DR^-^ area, 2) CD33^+^/IBA1^+^/HLA-DR^LOW^ area, and 3) CD33^+^ IBA1^+^ HLA-DR^HIGH^ area. From these areas, seven area fractions were calculated: 1) the CD33^+^ area of the total tumor area, 2) the area of MDSCs with no HLA-DR expression within the CD33^+^ area, 3) the area of MDSCs with low HLA-DR expression within the CD33^+^ area, 4) the total MDSC area, i.e., both HLA-DR^-^ and HLA-DR^LOW^, within the CD33^+^ area, 5) the area of MDSCs with no HLA-DR expression within the total tumor area, 6) the area of MDSCs with low HLA-DR expression within the total tumor area, and 7) the total MDSC area, i.e., both HLA-DR^-^ and HLA- DR^LOW^, within the total tumor area. All areas were analyzed for their association with survival and outlined in **Supplemental Figure 4**. A second algorithm was designed in the software module to quantify the entire HLA-DR+ area within the total tumor area (**Supplemental Figure 15**).

### CyTOF

Mass cytometry was performed in collaboration with the UCLA Jonsson Comprehensive Cancer Center (JCCC) and Center for AIDS Research Flow Cytometry Core Facility on a Fluidigm Helios CyTOF system. All antibodies were validated within the core, and those listed with heavy metal tags are listed in **Supplemental Figure 7** and determined to be non-overlapping by Maxpar Panel Designer Panel Wheel (Fluidigm). Cell were labeled with cisplatin (Cell-ID Cisplatin), a cocktail of metal-conjugated surface marker antibodies, and iridium (Cell-ID Intercalator) using reagents and protocols provided by Fluidigm (San Francisco, California, USA). Before analysis populations were cleaned by removing debris and dead cells before analysis **Supplemental Figure 8**. Samples were analyzed from six GBM patients at three timepoints for each patient (baseline, 2 months post-recurrence, and final sample collected). Three patients from this group had a good prognosis as denoted by survival >600 days post-resection and were still surviving, while three patients had a poor prognosis as denoted by a survival <600 days. Additionally, three LGG patients were analyzed at baseline for the comparison of baseline samples.

### CyTOF analysis

Prior to running CyTOF samples through data analysis, FCS files were normalized between runs using beads and the Nolan lab bead normalizer package [63]. The most current CyTOF data analysis tools were used for data analysis including multi-dimensional analysis with R following methods described by Nowicka et al. [39]. Additionally, FlowSOM analysis of CyTOF data was performed to identify changes in cell populations in an unbiased manner [43]. In a biased approach, CyTOF data was also analyzed using FlowJo software (Tree Star Inc.) as outlined in **Supplemental Figure 7**.

### Cytokine analysis

Cytokine analysis of patient samples was performed using a flow cytometry-based 65-plex cytokine array (Eve Technologies, Calgary, AB, Canada) **Supplemental Figure 11**.

### Statistical analyses

R version 3.4.4 was used in data analyses. The R function lm() was used to model cell percentages of lives cells as linear combinations of clinical covariates; as all values of such percentages were below 15%, saturation of percent was not a concern. The R functions survdiff() and coxph() of the R package survival were used to compute log-rank test P values and Cox proportional hazard model parameter P values, respectively.

## Author contributions

TJA, AGA, MAV and JDL provided conceptualization and design; TJA, AGA, MDS, DB, JV, ES, EEM-H, MS, JSH, MM, PH, MMG, and CAW performed the experiments; TJA, AGA, MDS, JV, TR, HIK, BWK, MAV, and JDL analyzed the data; TJA, DB, EEM-H, MAV, and JDL wrote the manuscript; HIK, BJW, MAV, and JDL provided financial support; and all authors provided final approval of the manuscript.

## Acknowledgements

We thank the patients treated at the Rose Ella Burkhardt Brain Tumor and Neuro-Oncology Center for donation of their blood samples for this study. We thank the staff of the Rose Ella Burkhardt Brain Tumor and Neuro-Oncology center for their collaboration acquiring samples. We thank the members of the Lathia laboratory and Dr. Ofer Reizes and his laboratory for insightful discussion and constructive comments on the manuscript. We thank Joseph Gerow and Eric Schultz for flow cytometry assistance. We thank the Janis V. Giorgi Flow Cytometry Core Laboratory, UCLA, for their assistance with CyTOF experiments. This work was funded by a National Institutes of Health grant (F31 NS101771, TJA), the Sontag Foundation (JDL), Blast GBM (JDL, MAV), the Cleveland Clinic VeloSano Bike Race (JDL, MAV), B*CURED (JDL, MAV), and Case Comprehensive Cancer Center (JDL, MAV).

